# Lecture-based, problem-based, digital problem-based and distance learning on knowledge improvement in medical education: a meta-analysis

**DOI:** 10.1101/2021.05.26.445870

**Authors:** Jiangming Sun

## Abstract

Problem-based learning (PBL), an educational approach well applied in education, was believed as a deep method that can promote problem solving, and critical thinking. Varies implementation of PBL across different settings were introduced. How to objectively evaluate knowledge effectiveness of PBLs remains a challenge. The present study is aiming to systematically investigate the knowledge improvement between types of PBL in medical education. Our meta-analysis showed that distance learning using digital PBL could be a good alternative to traditional learning in medical education.

## Background

Stimulating the development of learning skills such as problem solving, and critical thinking is a crucial goal of higher education nowadays^1^. Problem-based learning (PBL), an educational approach well applied in health professions education, was believed as a deep method that can promote clinical medical students to cultivate the actual clinical problems solving skills. Since PBL was introduced, implementation of PBL varies across different settings, such as digital problem-based learning (DPBL). Particularly, DPBL in the form of distance learning became more and more popular due to COVID-19 pandemic ^2, 3^. However, PBL is not universally. For example, lecture-based learning (LBL) is still the most commonly form in medical education in China^4^. Though, the hybrid PBL and LBL pedagogy or purely PBL approaches have been popular in China^4, 5^.

How to objectively evaluate knowledge effectiveness of various PBL remains a challenge. Luckily, certain randomized controlled trials (RCT) have been performed. On top of that, a few meta-analyses examined mean differences between types of learning. Here, this study is aiming to systematically investigate the knowledge improvement between types of PBL in medical education.

## Methods

### Study selection

Aiming for an objective evaluation, only randomized controlled trials (RCT) on LBL, PBL, hybrid LBL and PBL and DPBL in higher medical education passed quality assessment in previous meta-analysis^4-6^ were included in this analysis.

### Statistical analysis

Standardized mean difference (SMD) in knowledge outcome was calculated based on the available data from the included studies. 95% confidence interval of SMD was estimated using a R package MBESS^7^.

In current study, I assumed that there was one true effect size that underlined all the studies in the analysis. An inverse variance fixed-effect model was implemented consequently. A Student’s t-test was applied to examine differences in SMD from various comparisons.

## Results

### Study Characteristics

In total, 36 studies including 3969 subjects were examined in this work. Out of them, 26 were theory courses and 10 were practice courses. The later was mainly in the form of hybrid PBL and LBL except 1 PBL. Distribution of field of study was shown in Figure 1.

**Figure 1.**
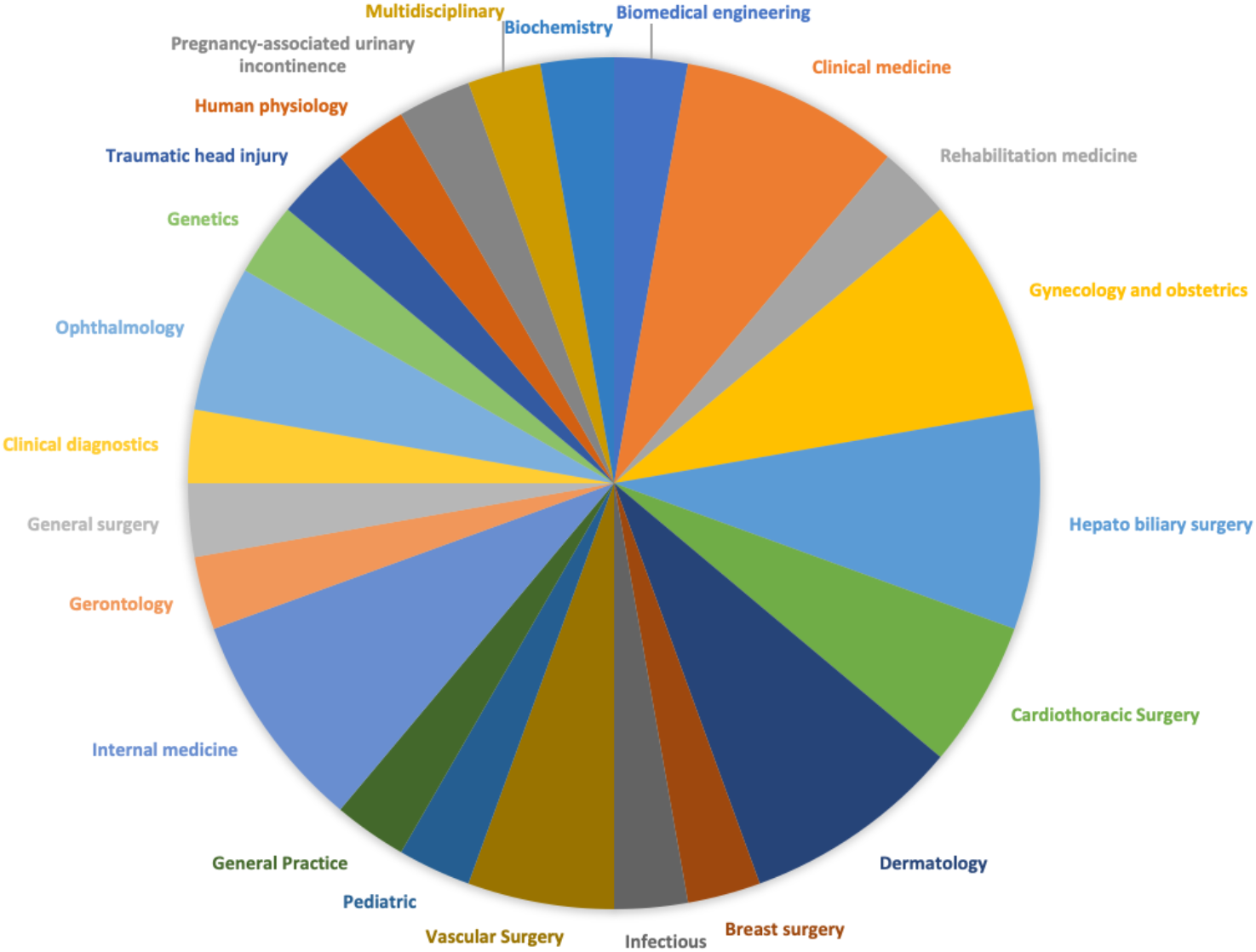
Pie chart showing field of study compositions.

### Effects of PBL, PBL-LBL hybrid or DPBL versus traditional LBL

As shown in Table 1, postintervention knowledge scores (SMD and 95% CI) were reported. Overall, PBL or a hybrid of PBL and LBL improved SMD greatly compared to LBL. Only 5 out of 28 studies failed to achieve a significant improvement.

**Table 1.**
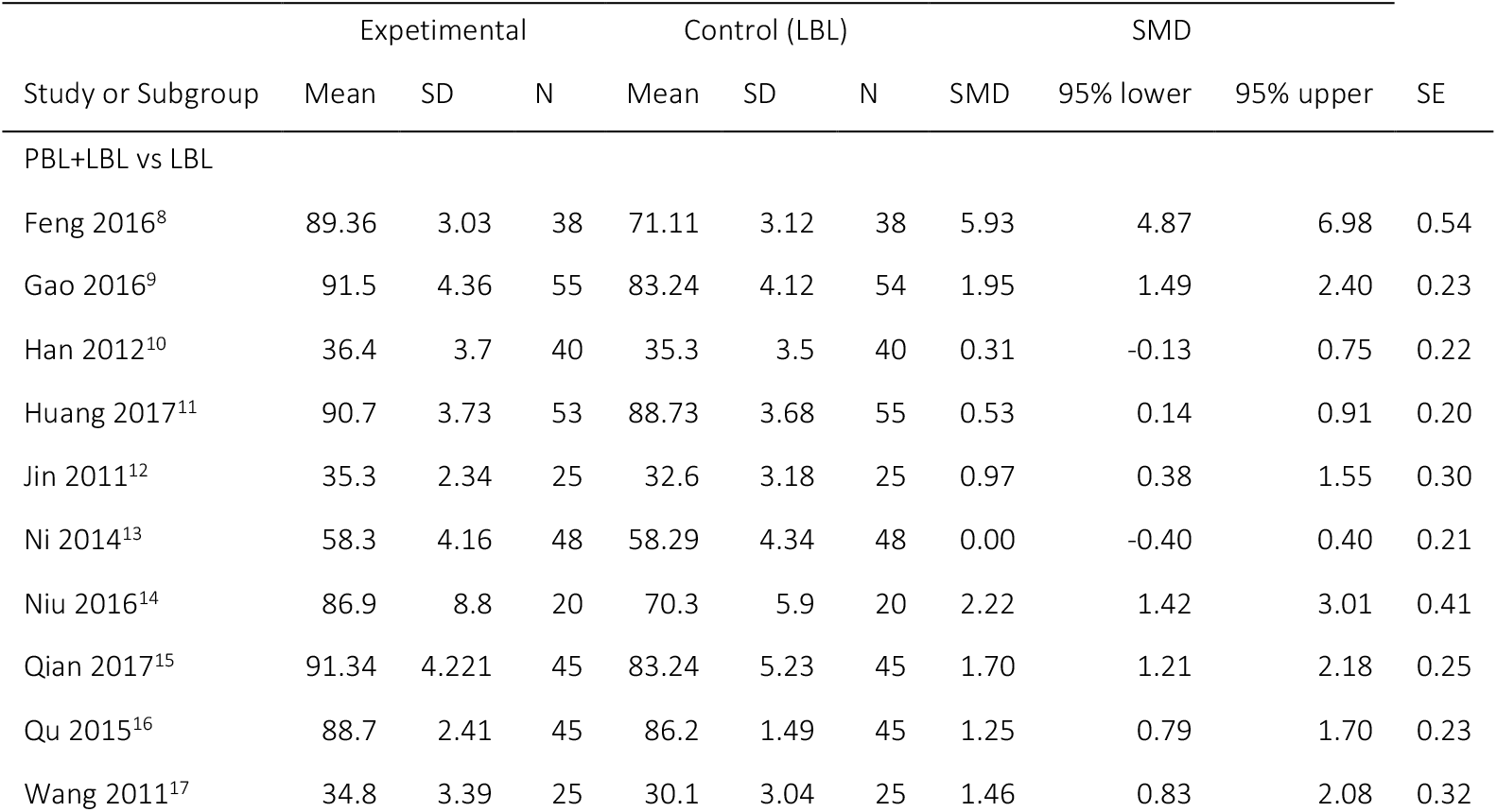
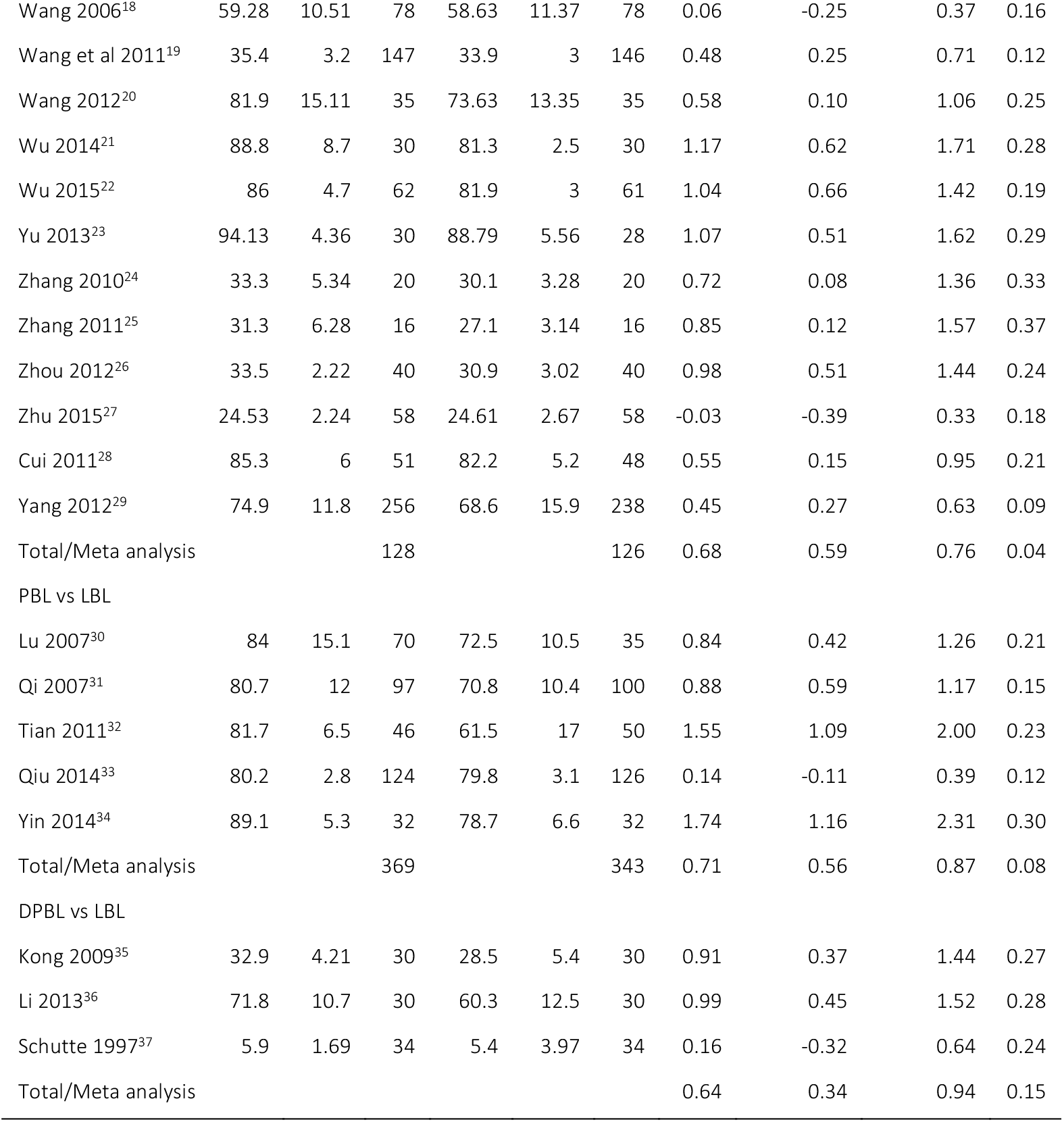
Knowledge outcomes from PBL+LBL, PBL, DPBL and traditional LBL.

Types of learning achieved greatest SMD versus traditional LBL was also examined (Figure 2). Among those 3 types of learning, PBL vs LBL achieved a highest SMD of 0.71 (95% CI 0.56-0.87), outperform PBL+LBL vs LBL (p=5.8 × 10^−21^) and DPBL vs LBL (p=0.046). No significant difference was found between PBL+LBL vs LBL and DPBL vs LBL.

**Figure 2.**
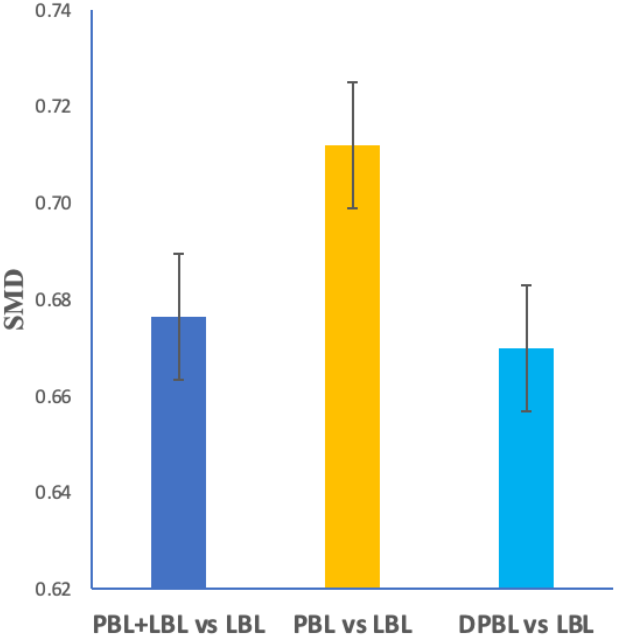
The pooled SMD using LBL as reference from the meta-analysis.

### Effects of various forms of DPBLs versus traditional PBL

The effects for comparisons between various settings of PBLs vs traditional PBL were retrieved from a study^6^. The pooled effects were given in Figure 3. No significant differences were found between DPBL and traditional PBL, as well as between partially distanced-based DPBL and PBL. However, a significant trend of knowledge improvements was found in DPBL vs PBL or partially distanced-based DPBL vs PBL, respectively.

**Figure 3.**
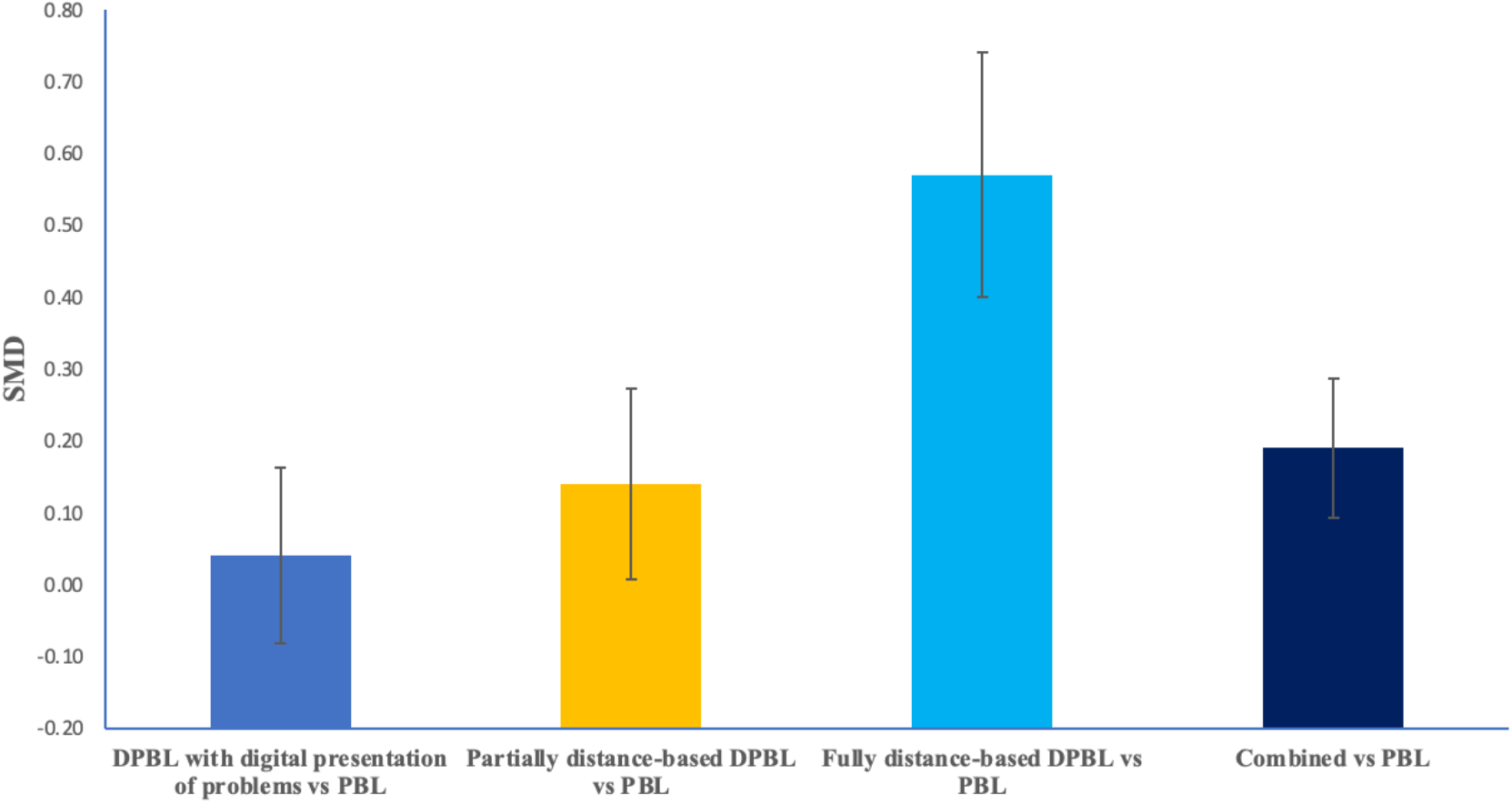
The pooled SMD using traditional PBL as reference from the meta-analysis. Summary statistics were retrieved from a study^6^.

The fully distance-based DPBL vs PBL achieved the highest knowledge outcome (SMD=0.57, 95% CI 0.23-0.92), greatly outperform partially distance-based learning DPBL vs PBL (p=4.6 × 10^−95^) and DPBL vs PBL (p=7.8 × 10^−78^). The partially distance-based learning DPBL vs PBL achieved a higher SMD than DPBL vs PBL (p=8.3 × 10^−18^). If all effects of DPBL vs PBL were merged, the SMD for combined effect vs PBL was 0.19 (95% CI 0-0.38).

## Discussion

The present study performed a meta-analysis on 36 RCT studies on PBL. As expected, PBL improved knowledge outcome greatly compared to LBL in higher medical education. Digital PBL was effective as a hybrid approach of PBL and LBL in comparison to LBL. Notably, fully distance based DPBL showed better knowledge scores in comparison to traditional PBL. This suggested that distance learning using digital PBL could be a good alternative to traditional learning in medical education which was certainly good news for education during this pandemic period of COVID-19 and could be future direction.

The meta-analysis was based on RCT studies which was quite rigorous and less likely be biased. Thus, sex difference should have less effect on conclusion from this study. One possible limitation was that most RCT studies on PBL and LBL were conducted in east Asian which might not correctly estimate true effect of knowledge improvement in other ethnic groups. However, the comparisons between partially or fully distance based DPBL and PBL employed participants diversely. As data on distance based DPBL is growing from last year, further investigations are hopefully an interesting next to evaluate its effectiveness in large scale.

